# Genome assembly and characterization of two Brazilian soybean cultivars contrasting for stink bug resistance

**DOI:** 10.64898/2026.11.06.764844

**Authors:** Marcos Antonio de Godoy Filho, João Paulo Gomes Viana, Lucas Borges dos Santos, Wei Wei, Michelle da Fonseca Santos, José Baldin Pinheiro

## Abstract

Brazil’s tropical soybean germplasm still has few genomic resources: only one assembly of a Brazilian cultivar is available at NCBI. Here we describe chromosome-level *de novo* assemblies of two contrasting cultivars: IAC-100, described as resistant to phytophagous stink bugs, and CD-215, susceptible. We sequenced high molecular weight DNA using 10x Genomics linked reads, assembled the reads with Supernova, anchored the assembly to the Williams 82 reference with RaGOO and corrected structural errors with REAPR. The assemblies span 1.057 and 1.061 Gb, with scaffold N50 values of 53.6 and 54.2 Mb (48.0 and 47.7 Mb excluding gaps), and recover more than 97% of conserved plant genes assessed by BUSCO. Homology-based projection of the reference protein-coding genes generated 46,774 and 46,620 gene models. Comparing the proteomes of the two cultivars, we identified 2,002 divergent isoforms in 1,427 loci, classified by sequence divergence, length variation, combined divergence and non-conserved homology. These data expand the genomic resources for tropical soybean germplasm, still underrepresented in public databases, and for studies of insect resistance.

## Background & Summary

Next-generation sequencing (NGS), dominated by short-read platforms such as Illumina, has reduced costs and made genomic data accessible on a large scale (Mardis 2017; Goodwin et al. 2016). However, in large and repetitive genomes, such as those of plants, short reads do not resolve repetitive regions, and *de novo* assemblies are fragmented into thousands of contigs, which limits the study of synteny, structural variants and gene families (Rice and Green 2019). Long reads from Pacific Biosciences (PacBio) and Oxford Nanopore (ONT) span these regions and generate much more contiguous assemblies (Jiao and Schneeberger 2017). Combined with chromosomal proximity mapping by Hi-C, which orders and orients contigs into chromosome-sized scaffolds (Lieberman-Aiden et al. 2009; Dudchenko et al. 2017), they have become the leading approach for reference genome assembly (Jung et al. 2019).

Linked reads offer a technological alternative for adding long-range information to short reads. High molecular weight DNA is partitioned into millions of compartments, and the fragments in each compartment receive the same barcode (Zheng et al. 2016). Reads that share a barcode are then grouped as originating from the same long molecule, which helps resolve repeats and build long scaffolds (Armstrong et al. 2019). The technology has been used for complex genomes of plants, such as those of pepper (*Capsicum annuum*; Hulse-Kemp et al. 2018) and pea (*Pisum sativum*; Kreplak et al. 2019), and of animals (Armstrong et al. 2019; Ozerov et al. 2018; Gao et al. 2023). The pioneering platform, 10x Genomics Chromium, has been discontinued for linked-read sequencing, but its principle is used in successor technologies, such as stLFR (Single-Tube Long Fragment Read) from MGI (Wang et al. 2019) and TELL-seq (Transposase Enzyme Linked Long-Read Sequencing) from UST (Chen et al. 2020). Both perform barcoding in a single tube, without complex microfluidics. stLFR uses co-barcoding on microbeads, and TELL-seq uses a transposase-based method. Because raw assemblies from any technology may contain structural errors, such as chimeras or collapsed regions, they are usually refined by reference-guided scaffolding (e.g., RaGOO; Alonge et al. 2019) and by error correction based on read remapping (e.g., REAPR; Hunt et al. 2013).

Soybean (*Glycine max* [L.] Merrill) has a large (*∼*1 Gbp), paleopolyploid genome whose repeat content is representative of the complexity encountered in many other crops (Schmutz et al. 2010). Despite its global economic importance, the available genomic resources are predominantly based on North American and Chinese germplasm (e.g., Williams 82, Zhonghuang 13; Schmutz et al. 2010; Shen et al. 2019). This gap is particularly critical for Brazil, the world’s largest producer, where the local germplasm, adapted to tropical conditions and a source of alleles for stress resistance, remains genomically and structurally underrepresented, with only one genome available at NCBI, published in 2021 by EMBRAPA (GenBank GCA_012273815.2). The absence of high-quality reference genomes for Brazilian cultivars represents a significant bottleneck for the advancement of precision plant breeding. Accessions adapted to equatorial and tropical latitudes are also underrepresented in existing global soybean pan-genomes (Liu et al. 2020; Bayer et al. 2022; Torkamaneh et al. 2021). More chromosome-level *de novo* assemblies of commercial cultivars and tropical local accessions are therefore needed, as well as the inclusion of this germplasm in public pan-genomes.

Phytophagous stink bugs (Hemiptera: Pentatomidae), such as *Euschistus heros* and *Nezara viridula*, are among the main soybean pests in Brazil and reduce seed yield and quality (Grolli et al. 2026). The cultivar IAC-100 is described in the literature as resistant to stink bugs, and has been used as a source of resistance in breeding populations, while CD-215 is susceptible (Rocha et al. 2015; Da Fonseca Santos et al. 2018; Grolli et al. 2026). However, the structural and sequence differences behind this contrast are not yet known, in part due to the lack of reference genomes for tropical genotypes. Here we describe chromosome-level *de novo* genome assemblies for both contrasting genotypes, the gene annotation projected onto them, the derived proteomes, the gene divergence classification between the two cultivars and the functional annotation (Gene Ontology and Pfam) of IAC-100. These assemblies serve as references for soybean germplasm grown in Brazil and can be used to develop molecular markers, map quantitative trait loci (QTLs) of agronomic interest and carry out comparative transcriptome studies. In addition, these assemblies can reduce the reference bias that arises when reads from tropical cultivars are mapped against Williams 82, a temperate-climate cultivar of distinct genetic origin, which lowers read mapping and the detection of variants in regions that differ from the reference. Together with transcriptome data, phenotyping or association mapping, the divergent loci described here can support future studies on the genetic basis of stink bug resistance in soybean.

## Methods

### DNA extraction, library construction and sequencing

High molecular weight (HMW) genomic DNA was extracted from young leaf tissue of the soybean cultivars *Glycine max* IAC-100 and CD-215, according to the protocol of Doyle and Doyle (1987). The integrity of the extracted DNA was assessed by gel electrophoresis to confirm the presence of long fragments.

Linked-read libraries were prepared with the Chromium Genome & Gel Bead Kit and processed on the 10x Genomics Chromium Controller (10x Genomics, Pleasanton, CA, USA), according to the manufacturer’s specifications. The libraries were quantified and sequenced on the Illumina NovaSeq 6000 platform to generate 2 *×* 150-bp paired-end reads.

### Genome assembly and refinement pipeline

#### *De novo* assembly from linked reads

The quality of raw sequencing data was assessed with FastQC (v0.11.9; Andrews 2010). *De novo* assembly was performed with Supernova (v2.1.1; Weisenfeld et al. 2017), which employs a de Bruijn graph algorithm optimized for linked-read data. As recommended by 10x Genomics, the reads were subsampled with the --maxreads parameter to a target coverage of about 56*×*, preventing the introduction of assembly artifacts from an excess of accumulated sequencing errors. The output in pseudohap format, which provides a haploid representation of the assembly, was used for all subsequent steps.

#### Reference-guided scaffolding

The Supernova contigs were ordered and oriented into pseudomolecules using RaGOO (v1.1; Alonge et al. 2019). The reference genome of the cultivar *Glycine max* Williams 82 (RefSeq GCF_000004515.5) was used to guide the process. RaGOO employs Minimap2 (v2.17; Li 2018) to align contigs against the reference and infer their chromosomal position and orientation.

#### Error correction and iterative refinement

Structural errors in the pseudomolecules were corrected with REAPR (v1.0.17; Hunt et al. 2013). Illumina reads trimmed with Trimmomatic (Bolger et al. 2014) were mapped back to the scaffolds using BWA (v0.7.17; Li and Durbin 2009). REAPR analyzed the mappings to identify regions with error signatures and broke the scaffolds at the detected points. The REAPR alerts were grouped into two categories: (1) positional uncertainty zones, with the alerts *Low fragment coverage within a contig* and *Low fragment coverage over a gap*, which indicate regions with low read coverage, and (2) structural assembly errors, with the alerts *FCD (Fragment Coverage Distribution) error within a contig* and *over a gap*, which point to assembly errors (misassemblies) based on deviations in the fragment size distribution and on discordant read pairs.

The corrected contigs were then scaffolded again with RaGOO (Alonge et al. 2019), with the same parameters as before, to produce the final assemblies.

### Assembly quality assessment and annotation

#### Completeness and contiguity analysis

The gene content completeness of the assemblies and of the Williams 82 reference was assessed by BUSCO (v3.0.2; Waterhouse et al. 2018) against the embryophyte ortholog database (embryophyta_odb10; Kriventseva et al. 2019). Assembly contiguity statistics, including scaffold N50 and the number of scaffolds containing 75% of the assembly, were calculated to quantify structural quality, both with gap (N) positions included in the sequence length and with them excluded.

#### Structural annotation based on homologous projection

Repetitive elements were first identified and masked with RepeatMasker (v4.1.1; Smit et al. 2013–2015) with the SoyTEdb (Du et al. 2010) and Dfam (v3.1; Hubley et al. 2016; Storer et al. 2021) libraries. Then, to make the annotation comparable between the assemblies, the genes of Williams 82 (RefSeq GCF_000004515.5) were projected onto the IAC-100 and CD-215 assemblies. The reference GFF3 annotation file was first filtered with gffread (v0.12.7; Pertea and Pertea 2020) using the -C parameter, retaining only transcripts containing a valid coding sequence (CDS). The projection was done with Liftoff (v1.6.3; Shumate and Salzberg 2021), which aligns the reference features with Minimap2 (v2.28-r1209; Li 2018), with the -copies parameter, which also maps gene duplications and paralogs. The corresponding amino acid sequences (proteomes in FASTA format) were extracted using gffread.

### Comparative genomics and gene divergence classification

To directly compare gene content between the assembled genomes and classify variations at the protein level, a comparative genomics pipeline was implemented, divided into three consecutive steps based on the predicted proteome:

(i) Gene pairing: Each IAC-100 isoform was paired with the CD-215 isoform projected from the same Williams 82 transcript in the same genomic region. Orthogroups were also inferred with OrthoFinder (v2.5.5; Emms and Kelly 2019) and used in Technical Validation II to check the pairing.
(ii) Isoforms without projection: IAC-100 isoforms not projected onto CD-215 were searched with TBLASTN (BLAST+ v2.17.0; Camacho et al. 2009; *E ≤* 10*^−^*^10^) against the CD-215 assembly. Those whose best hit had *<*80% identity or *<*50% coverage were classified as Non-Conserved Homologs (NCH), provided that TBLASTN found the protein in its own IAC-100 gene and that the matching CD-215 region was not an assembly gap.
(iii) Homolog Divergence: For each pair, proteins were cut at the first stop codon and aligned end to end with the Parasail library (Daily 2016; BLOSUM62, Henikoff and Henikoff 1992; gap penalties 10/1). Based on similarity (*≥*98%) and alignment coverage (*≥*95%) thresholds, with coverage calculated on the longer of the two aligned sequences to neutralize asymmetry biases, isoforms were classified into four mutually exclusive categories: Conserved (CON; coverage *≥*95% and similarity *≥*98%), Sequence Divergence (SDV; coverage *≥*95% and similarity *<*98%), Structural Divergence (STV; coverage *<*95% and similarity *≥*98%) and Combined Divergence (CDV; coverage *<*95% and similarity *<*98%). Classes that could be caused by assembly gaps (N) were not assigned.

### Functional annotation and Gene Ontology enrichment

The assignment of Gene Ontology (GO) terms (Ashburner et al. 2000; The Gene Ontology Consortium 2026) to the whole IAC-100 proteome was performed *de novo*, combining two independent methodologies. The sequences were submitted to InterProScan (v5.59-91.0; Jones et al. 2014) for the identification of structural domains and conserved motifs, querying the Pfam (v35.0; Mistry et al. 2021) and PANTHER (v17.0; Thomas et al. 2022) databases. In parallel, orthologous mapping was conducted with eggNOG-mapper (v2.1.13; Cantalapiedra et al. 2021) and the eggNOG database (v5.0.2; Huerta-Cepas et al. 2019). GO terms inferred by either tool were consolidated per locus, as the union of the terms of all its isoforms, to ensure comprehensive functional characterization. The consolidated GO annotation was curated to remove obsolete terms, replaced by their declared successors where available, and terms not applicable to plants. GO terms were then propagated to their ancestors through the *is_a* and *part_of* relations of the ontology.

Statistical over-representation analyses (GO enrichment) were conducted using a one-sided Fisher’s Exact Test. To ensure statistical validity, the analysis universe was strictly restricted to the loci of the five classes (CON, SDV, STV, CDV and NCH) with at least one GO term assigned following the functional annotation step (annotated background = 31,772 loci). The same annotation was used for the divergence classes and for the background. Genes without GO terms were excluded from both the background and the target groups prior to the construction of 2 *×* 2 contingency tables, as their inclusion would artificially inflate the true negatives, following standard practice in genomic enrichment analyses (Rivals et al. 2007). Only terms assigned to 10 to 500 background loci and to at least one locus of the class were tested. False discovery rate (FDR) values were calculated by the Benjamini-Hochberg method (Benjamini and Hochberg 1995) within each class, with the three GO namespaces together, and terms with FDR *<* 0.05 were considered significant. The SDV, STV and CDV classes were tested. NCH (15 loci) was not tested because of its small size. Significant terms were interpreted through the genes that carry them.

For an overview, each term was associated with its level-2 ancestors in the Gene Ontology (the direct children of the three roots of the ontology), as in WEGO (Ye et al. 2018). For each tested class and for the background, the percentage of annotated loci carrying each level-2 term was computed without statistical testing.

## Data Records

The linked reads of IAC-100 and CD-215 are in the NCBI Sequence Read Archive under accessions SRP749030 and SRP749041 (BioProjects PRJNA1506754 and PRJNA1506755, BioSamples SAMN62148116 and SAMN62148178, respectively), as paired-end FASTQ files that retain the 10x Genomics barcode structure. The assemblies are in DDBJ/ENA/GenBank under accessions JCFFPV000000000 (IAC-100, assembly IAC-100_v1.0) and JCFFPU000000000 (CD-215, assembly CD-215_v1.0), in FASTA format, with the 20 pseudomolecules following the chromosome numbering of Williams 82 (IAC100_Gm01 to IAC100_Gm20 and CD215_Gm01 to CD215_Gm20, names also used in the GFF3 files). The versions described here are JCFFPV010000000 and JCFFPU010000000.

The assemblies (IAC_100.fasta.gz and CD_215.fasta.gz), the remaining files and the code are in a single Zenodo record (https://doi.org/10.5281/zenodo.23805779). The projected annotations are in IAC_100.v1.0_liftoff.gff3 and CD_215.v1.0_liftoff.gff3, with the coordinates on the corresponding assembly, the gene and transcript identifiers of the Williams 82 RefSeq annotation (e.g., gene-LOC102661143 and rna-XM_014762920.1) and the projection quality attributes written by Liftoff (e.g., coverage, sequence_ID, copy_num_ID, matches_ref_protein and valid_ORF). The proteomes are in IAC_100_ protein_liftoff.fasta and CD_215_protein_liftoff.fasta. In these sequences, X marks an internal stop codon or a codon with unknown bases. The divergence classification is in IAC_100_divergence_classes.tsv, with one row per classified IAC-100 isoform and the columns transcript_id, gene_id (locus) and class (CON, SDV, STV, CDV or NCH). The functional annotation of IAC-100 is in IAC_100_functional_annotation.tsv, with one row per locus with at least one GO term or Pfam domain and the columns gene_id, go_terms (curated GO terms, before propagation) and pfam_domains (Pfam domains of the isoforms of the locus), with the identifiers within a cell separated by semicolons. Both tables are tab-separated.

## Technical Validation

### Technical Validation I: Assembly contiguity and completeness

Sequencing of soybean cultivars IAC-100 and CD-215 using the 10x Genomics Chromium platform generated paired-end reads, which were subsampled to 460 million reads (230 million read pairs) per genome with --maxreads (Table 1). After filtering, the mean read length was 138.50 bp, resulting in an effective genome coverage of 44.25*×* for IAC-100 and 43.54*×* for CD-215.

**Table 1.** Descriptive statistics of the raw genome assemblies generated by Supernova. Total coverage and estimated genome size from Supernova.

| Parameters | IAC-100 | CD-215 |
|---|---|---|
| Number of reads used | 460.02 M | 460.02 M |
| Mean read length after filtering | 138.50 bp | 138.50 bp |
| Total coverage | 58.91× | 58.65× |
| Effective coverage | 44.25× | 43.54× |
| Estimated genome size | 1.17 Gb | 1.18 Gb |
| HMW DNA molecule length | 65.74 kb | 74.82 kb |
| Number of scaffolds | 31,518 | 31,455 |
| Number of scaffolds > 10,000 bp | 5,822 | 5,730 |
| Scaffold N50 | 3.69 Mb | 3.94 Mb |
| Scaffold N50 > 10,000 bp | 4.18 Mb | 4.58 Mb |
| Scaffold N50, gaps excluded (≥ 10,000 bp) | 4.02 Mb | 4.28 Mb |
| GC content | 34.56% | 34.52% |
| Assembly size | 1.052 Gb | 1.055 Gb |
| Assembly size > 10,000 bp | 959 Mb | 963 Mb |
| Number of scaffolds with 75% of assembly > 10,000 bp | 159 | 144 |

The metrics of the initial assembly are detailed in Table 1. The total assembly size was 1.052 Gb (IAC-100) and 1.055 Gb (CD-215), corresponding to approximately 90% (80% excluding gaps) of the estimated genome size (1.17 to 1.18 Gb). The contiguity of the raw assemblies, assessed by scaffold N50, was 3.69 Mb and 3.94 Mb for IAC-100 and CD-215, respectively. Considering only scaffolds larger than 10,000 bp, the N50 increased to 4.18 Mb and 4.58 Mb, or 4.02 Mb and 4.28 Mb when gap positions are excluded, as reported by Supernova. Notably, in scaffolds of at least 10,000 bp, 75% of the assembled sequence was contained in only 159 (IAC-100) and 144 (CD-215) scaffolds, indicating an initial assembly of high quality and contiguity.

This contiguity, obtained before refinement, indicates that the linked-read data are of good quality. The slightly greater contiguity of the initial assembly of CD-215 is probably due to the longer mean length of its DNA molecules (74.82 kb vs. 65.74 kb), which usually increases contiguity in linked-read assemblies (Armstrong et al. 2019).

The metrics obtained in the initial assembly step are consistent with results described for pepper (*Capsicum annuum*), whose scaffold N50 reached 3.69 Mb (Hulse-Kemp et al. 2018).

In plant genomes, the abundance of repeats limits assemblies made only with short reads. In contrast, genomes with fewer repeats, such as those of the European perch (*Perca fluviatilis*; N50 of 6.3 Mb; Ozerov et al. 2018) or the African wild dog (*Lycaon pictus*; N50 of up to 21 Mb; Armstrong et al. 2019), tend to achieve greater contiguity with similar approaches. The contiguity obtained here is therefore within what is expected for linked-read assemblies of repeat-rich plant genomes.

During refinement, REAPR identified 33,387 and 34,787 putative structural alerts in the IAC-100 and CD-215 genomes, respectively. Of these, 71.8% (IAC-100) and 74.0% (CD-215) were positional uncertainty zones, and 28.2% and 26.0% were structural errors of the *de novo* assembly (misassemblies), which REAPR corrected.

The refinement resulted in a substantial improvement in genome quality. As presented in Table 2, the final scaffold N50 reached 53.6 Mb for IAC-100 and 54.2 Mb for CD-215, or 48.0 Mb and 47.7 Mb when gap positions are excluded, representing an increase of approximately 11- to 12-fold (11.9*×* and 11.1*×*, respectively, excluding gaps) relative to the initial assembly. In each cultivar, 75% of the genome is now in 15 scaffolds. The total number of sequences was reduced to 2,846 (IAC-100) and 2,608 (CD-215): 20 chromosome-scale pseudomolecules, which hold 97.9% and 98.0% of the assembled length, and 2,826 and 2,588 unplaced sequences (22.4 and 21.0 Mb). The total assembly size is 1.057 Gb and 1.061 Gb, respectively, of which 11.5% and 12.0% are gap positions (N), leaving 935.4 and 933.5 Mb of assembled sequence.

**Table 2.** Comparison of assembly quality parameters for the generated assemblies and for the Williams 82 (GCF_000004515.5) and Zhonghuang 13 (GCA_003349995.2) cultivars, computed from the FASTA files.

| Metric | IAC-100 | CD-215 | Williams 82 | Zhonghuang 13 |
|---|---|---|---|---|
| Scaffold N50 | 53.6 Mb | 54.2 Mb | 48.58 Mb | 51.95 Mb |
| Scaffold N50, gaps excluded | 48.0 Mb | 47.7 Mb | 47.8 Mb | 51.8 Mb |
| Number of sequences | 2,846 | 2,608 | 1,192 | 45 |
| Assembly size | 1.057 Gb | 1.061 Gb | 979 Mb | 1.011 Gb |
| Assembly size, gaps excluded | 935.4 Mb | 933.5 Mb | 955.9 Mb | 1,009.0 Mb |
| Gap positions (N) | 11.5% | 12.0% | 2.4% | 0.2% |

The chromosome-scale contiguity of the final assemblies is comparable to that of the soybean references Williams 82 (48.58 Mb), the scaffolding reference of this study, and Zhonghuang 13 (51.95 Mb; Shen et al. 2019; Table 2). When gap positions are excluded, the scaffold N50 of IAC-100 and CD-215 (48.0 and 47.7 Mb) is essentially that of Williams 82 (47.8 Mb), as expected from reference-guided scaffolding, which reproduces the chromosome structure of the reference. With gaps included, the values (53.6 and 54.2 Mb) exceed those of the two references only because the IAC-100 and CD-215 assemblies contain a larger proportion of gap positions (11.5% and 12.0%, against 2.4% in Williams 82 and 0.2% in Zhonghuang 13). Excluding gaps, they also contain less sequence than the reference (935.4 and 933.5 Mb, against 955.9 Mb). The IAC-100 and CD-215 assemblies were obtained via scaffolding guided by the Williams 82 genome itself, a faster and lower-cost alternative to Hi-C, but one that limits the independent discovery of macro-structural rearrangements specific to these cultivars. The relevance of these assemblies lies in covering Brazilian tropical cultivars previously lacking structural genomic resources.

Figure 1 demonstrates that both assemblies are highly complete, with 97.1% (CD-215) and 97.6% (IAC-100) of BUSCO genes identified as complete. As in the Williams 82 reference genome (741 duplicated and 609 single-copy), more BUSCO genes were found in duplicated copies (754 in IAC-100 and 725 in CD-215) than in single copy (588 and 610), as expected for the paleopolyploid soybean genome.

**Figure 1.**
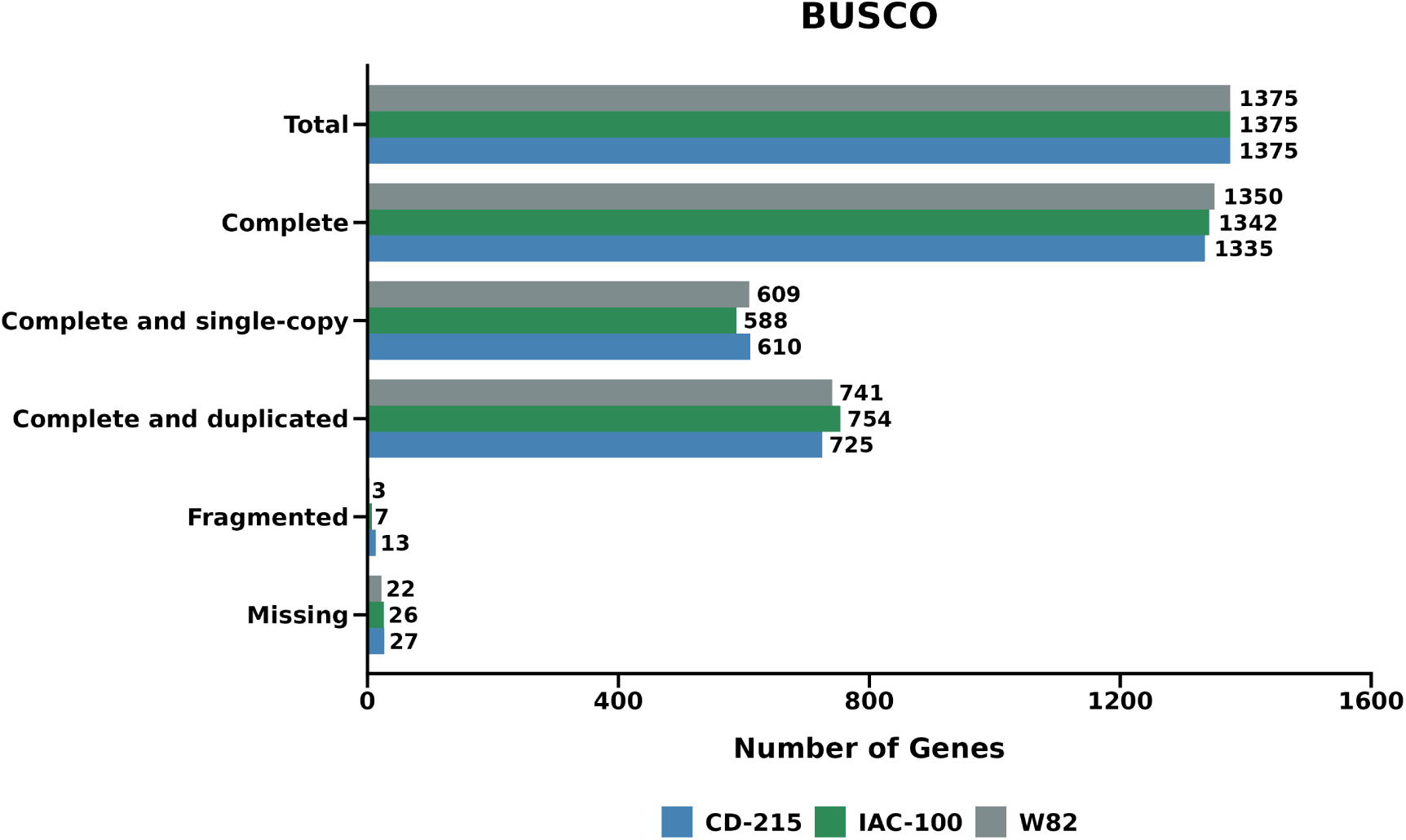
Results of the BUSCO (Benchmarking Universal Single-Copy Orthologs) analysis for the assembled genomes of IAC-100 and CD-215, compared with the Williams 82 reference genome (W82). The graph shows the number of complete (single-copy and duplicated), fragmented and missing orthologous genes.

In the structural annotation, the identification of repetitive elements with RepeatMasker indicated that approximately 44% of each assembled genome is composed of repetitive sequences, a proportion highly consistent with that observed in the Williams 82 reference genome (*∼*43%). Subsequently, homologous projection with Liftoff identified 46,774 and 46,620 gene models in the IAC-100 and CD-215 genomes, respectively, corresponding to 46,067 and 46,047 (98.4%) of the 46,794 protein-coding genes of the filtered reference, plus 707 and 573 additional copies detected with the -copies option. The set of 46,774 gene models projected for IAC-100 produced 70,736 protein sequences. This decrease in the number of predicted genes relative to the total from the original reference (from 59,668 gene and pseudogene loci of the reference annotation to *∼*47,000) is due to the gffread filter (-C), which eliminated pseudogenes and non-coding RNA genes. This gene set is the basis of the comparison between the cultivars.

### Technical Validation II: Annotation consistency and orthology

The structural annotation of the two cultivars was projected from Williams 82 with Liftoff. As a result, the gene order and the exon and intron boundaries are defined in the same way in both assemblies, which allows the proteins to be compared directly. Because it was projected from Williams 82, the annotation does not include genes present in the cultivars and absent from Williams 82.

Of the IAC-100 transcripts compared with those of CD-215, 61,987 of the 63,989 classified isoforms (96.9%) were Conserved (CON), indicating high protein conservation between the two cultivars. In contrast, we identified 2,002 divergent isoforms, with sequence or length differences between the resistant (IAC-100) and the susceptible (CD-215) cultivar. Because of alternative splicing, several of them belong to the same locus. To avoid overestimating the variation, the counts are given per isoform and per locus (Table 3). Together, the divergent isoforms map to 1,427 distinct loci. The per-class counts sum to more than this because a locus may carry isoforms in more than one class. As a check of the pairing, the proportion of pairs whose two isoforms fall in the same OrthoFinder orthogroup was 99.8% for CON, 89.9% for SDV, 61.8% for STV and 26.6% for CDV. This proportion falls as divergence increases because orthogroups are built from sequence similarity.

**Table 3.**
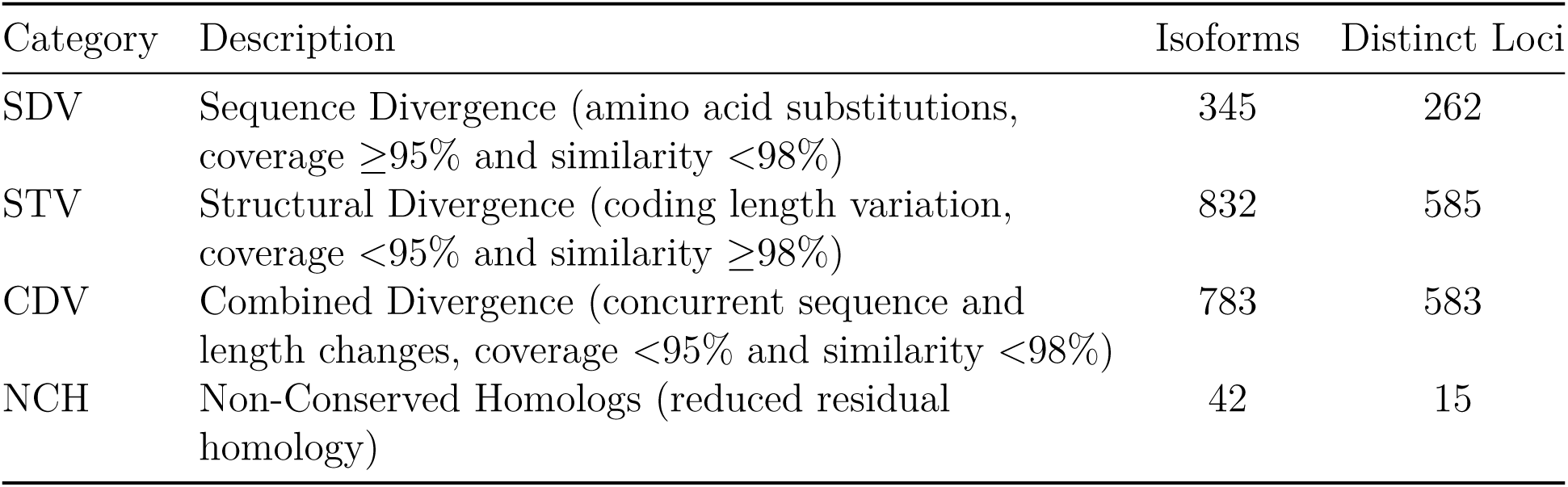
Divergence classes of IAC-100 relative to CD-215, in numbers of isoforms and of distinct loci. Distinct loci: isoforms collapsed to their parent gene in the projected annotation, with additional copies from -copies counted with the reference gene they derive from.

| Category | Description | Isoforms | Distinct Loci |
|---|---|---|---|
| SDV | Sequence Divergence (amino acid substitutions, coverage ≥95% and similarity <98%) | 345 | 262 |
| STV | Structural Divergence (coding length variation, coverage <95% and similarity ≥98%) | 832 | 585 |
| CDV | Combined Divergence (concurrent sequence and length changes, coverage <95% and similarity <98%) | 783 | 583 |
| NCH | Non-Conserved Homologs (reduced residual homology) | 42 | 15 |

### Technical Validation III: Functional coherence of the annotation

The functional annotation of the IAC-100 assembly was assessed by testing whether the Gene Ontology terms recovered for each tested divergence class are enriched relative to the background. Of the loci of each tested class, 170 of 262 (SDV), 402 of 585 (STV) and 317 of 583 (CDV) had at least one GO term and entered the test.

A single term was significant: ADP binding (GO:0043531), in SDV and in CDV. In SDV it was assigned to 12 of the 170 annotated loci (FDR = 1.2 *×* 10*^−^*^5^) and in CDV to 13 of the 317 (FDR = 3.1 *×* 10*^−^*^3^), against 252 of the 31,772 background loci. Of these loci (24 in total, one of them in both classes), 23 are annotated with an NB-ARC domain, the nucleotide-binding domain of nucleotide-binding leucine-rich repeat (NLR) disease resistance proteins, and they include Rj2 (SDV). No term was significant in STV, where the same term had the lowest *P* value (FDR = 0.17).

In a descriptive count of the Pfam domains assigned by InterProScan (Table 4), each tested class has 12 loci with an NB-ARC domain, or 6.8% (SDV), 2.8% (STV) and 3.6% (CDV) of its loci with a Pfam domain, against 0.6% of the background. In total, there are 34 distinct loci, all of which carry the ADP binding term. Domains of transposable-element proteins, such as transposase, reverse transcriptase, integrase and gag, occur in 21.5% (SDV), 16.7% (STV) and 13.3% (CDV) of the loci with a Pfam domain, against 3.6% of the background, whereas the protein kinase and pentatricopeptide repeat (PPR) domains, which belong to large gene families, occur in 3.4 to 5.6% and 2.1 to 2.8% of these loci, respectively, against 5.3% and 1.8% of the background.

**Table 4.** Loci carrying selected Pfam domains in the SDV, STV and CDV classes and in the background (loci of the five classes). Each cell gives the number of loci with the domain, the number of loci with at least one Pfam domain and the percentage, without statistical testing. Domains: NB-ARC (PF00931), protein kinase (PF00069 or PF07714), PPR (PF01535 or PF13041) and the 28 transposable-element entries listed in te_pfam.tsv, provided with the code.

| Domain | SDV | STV | CDV | Background |
|---|---|---|---|---|
| NB-ARC domain | 12 / 177 (6.8%) | 12 / 430 (2.8%) | 12 / 331 (3.6%) | 202 / 34,262 (0.6%) |
| Transposable-element domains | 38 / 177 (21.5%) | 72 / 430 (16.7%) | 44 / 331 (13.3%) | 1,232 / 34,262 (3.6%) |
| Protein kinase domain | 6 / 177 (3.4%) | 24 / 430 (5.6%) | 13 / 331 (3.9%) | 1,805 / 34,262 (5.3%) |
| PPR repeat | 5 / 177 (2.8%) | 9 / 430 (2.1%) | 9 / 331 (2.7%) | 631 / 34,262 (1.8%) |

Across the level-2 GO terms, the three classes have the same four predominant terms as the background: cellular process, cellular anatomical structure, binding and catalytic activity (Figure 2).

**Figure 2.**
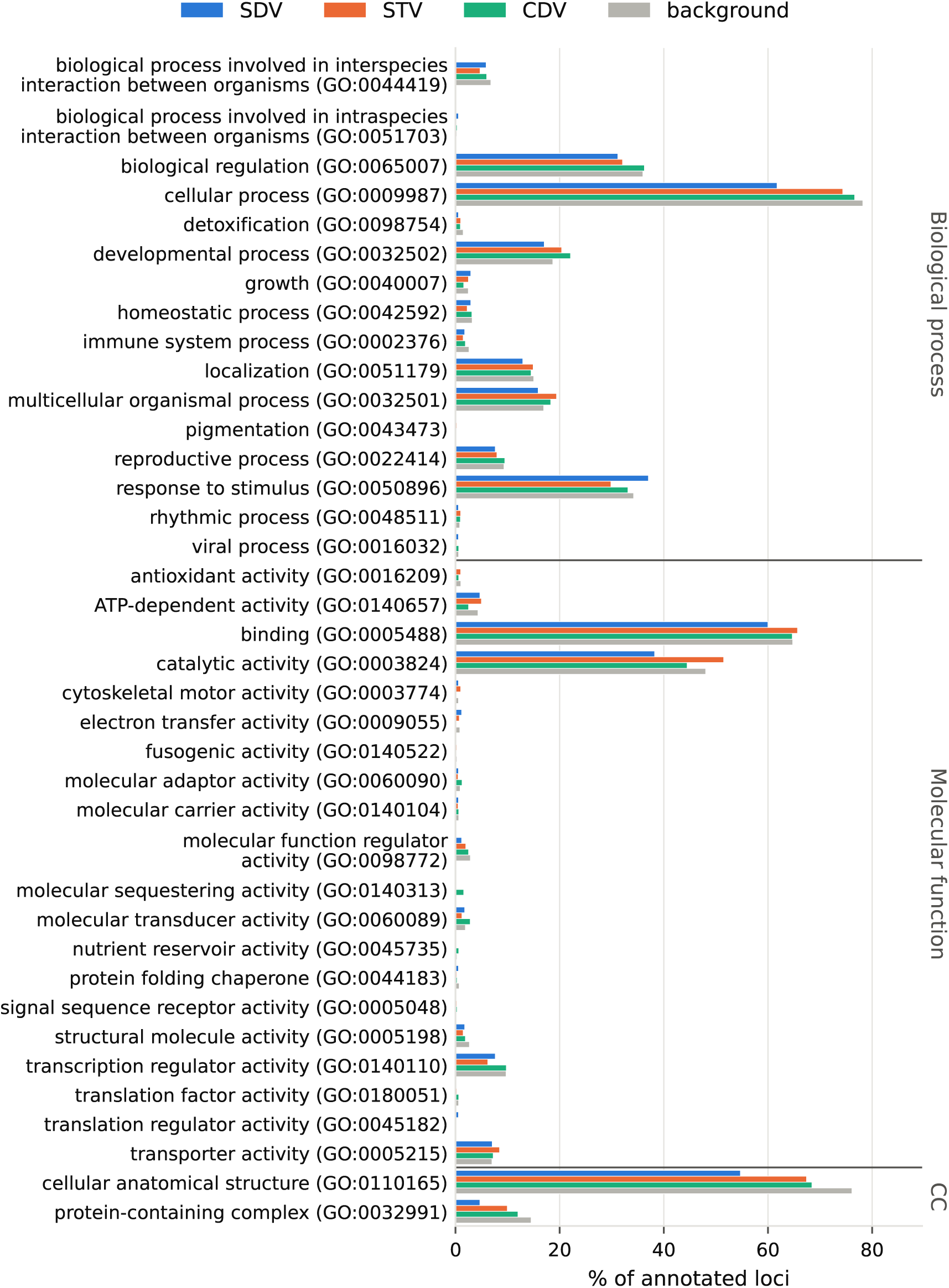
Level-2 GO terms by divergence class. The terms are grouped by ontology (biological process, molecular function and cellular component, CC). The X-axis shows the percentage of annotated loci in each group (SDV, STV, CDV and background) that carry the term, which allows groups of different sizes to be compared.

These differences (SDV, STV and CDV) are natural variation in soybean germplasm. Although IAC-100 is described as resistant to stink bugs, the divergences found here mainly reflect the different breeding histories, the crosses and the genetic drift of the two cultivars. Therefore, a divergence does not, by itself, indicate a relationship with insect defense.

The divergence classification generated constitutes a genetic resource map for the tropical germplasm, in which the resistance determinants represent a subset that needs to be delimited by future quantitative genetics studies (e.g., QTL mapping or genome-wide association studies, GWAS) and functional validation. The coordinates of the divergent loci can support comparative genomics and the development of molecular markers for agronomic traits.

## Data Availability

The raw reads are in the NCBI Sequence Read Archive under accessions SRP749030 (IAC-100, BioProject PRJNA1506754) and SRP749041 (CD-215, BioProject PRJNA1506755), and the assemblies are in DDBJ/ENA/GenBank under JCFFPV000000000 (IAC-100) and JCFFPU000000000 (CD-215). The assemblies, the annotations, the proteomes, the divergence classification, the functional annotation and the code are in Zenodo (https://doi.org/10.5281/zenodo.23805779).

## Code Availability

The scripts used to project the annotation, classify gene divergence and perform the Gene Ontology and Pfam analyses are in the same Zenodo record as the data (https://doi.org/10.5281/zenodo.23805779). The analyses were performed in Python 3.13, with biopython v1.87 (Cock et al. 2009), parasail v1.3.4, scipy v1.17.1 (Virtanen et al. 2020) and matplotlib v3.10.9 (Hunter 2007).

## Funding

This work was supported by the Coordenação de Aperfeiçoamento de Pessoal de Nível Superior (CAPES) and the Conselho Nacional de Desenvolvimento Científico e Tecnológico (CNPq).

## Notes

https://doi.org/10.5281/zenodo.23805779

https://www.ncbi.nlm.nih.gov/bioproject/PRJNA1506754

https://www.ncbi.nlm.nih.gov/bioproject/PRJNA1506755

### Competing Interest Statement

The authors have declared no competing interest.

